# Species-specific design of artificial promoters by transfer-learning based generative deep-learning model

**DOI:** 10.1101/2023.12.27.573416

**Authors:** Yan Xia, Xiaowen Du, Bin Liu, Shuyuan Guo, Yi-Xin Huo

## Abstract

Native prokaryotic promoters share common sequence patterns, but are species dependent. For understudied species with limited data, it is challenging to predict the strength of existing promoters and generate novel promoters. Here, we developed PromoGen, a collection of nucleotide language models to generate species-specific functional promoters, across dozens of species in a data and parameter efficient way. Twenty-seven species-specific models in this collection were finetuned from the pretrained model which was trained on multi-species promoters. When systematically compared with native promoters, the *Escherichia coli-* and *Bacillus subtilis-*specific artificial PromoGen-generated promoters (PGPs) were demonstrated to hold all distribution patterns of native promoters. A regression model was developed to score generated either by PromoGen or by another competitive neural network, and the overall score of PGPs is higher. Encouraged by *in silico* analysis, we further experimentally characterized twenty-two *B. subtilis* PGPs, results showed that four of tested PGPs reached the strong promoter level while all were active. Furthermore, we developed a user-friendly website to generate species-specific promoters for 27 different species by PromoGen. This work presented an efficient deep-learning strategy for *de novo* species-specific promoter generation even with limited datasets, providing valuable promoter toolboxes especially for the metabolic engineering of understudied microorganisms.

## Introduction

Regulating gene expression is essential to metabolic and biosynthetic engineering, as it enables the optimization of the production of desired metabolites, biofuels, pharmaceuticals, and other valuable compounds (1–4). Promoters are core regulatory elements that initiate the gene transcription, regulate gene expression, and affect metabolic flux distribution in metabolic pathways (5,6). By obtaining well-understood and user-friendly promoters, scientists could redesign cells to create gene circuits and synthetic biological systems tailoring to specific requirements (7). Although have been utilized to regulate gene expressions, native promoters lack continuous regulatory strength and wide regulatory scope (8). Therefore, many efforts have been made to develop artificial or hybrid promoters for regulating gene expressions (9–11).

The field of promoter engineering utilized random and semi-rational mutagenesis techniques to develop promoter libraries (12,13) that containing promoters with varied expression levels. For example, error-prone PCR based random mutagenesis was applied to the PL-λ promoter in *Escherichia coli*, leading to a substantial 196-fold variation in promoter activity identified through high-throughput screening (14). This approach has also been successfully applied to other organisms, including *Lactococcus lactis*, *Pichia pastoris*, and *Saccharomyces cerevisiae* Tef1 to generate active promoters (15–17). On the other hand, semi-rational design has been used to engineer *Bacillus subtilis* synthetic promoter libraries to obtain promoters with transcriptional activities varying up to 100-fold by fixing the −10 and −35 regions and altering the flanking regions of promoters *P_veg_*, *P_ser_*_A_, and *P_ymd_*_A_ (16,18). Although engineered promoters with enhanced performance have been reported, challenges remained in generating and screening large-scale libraries for obtaining promoters with a broad and continuous range of strengths (8).

Deep learning architectures have shown substantial promise for the manipulation of both eukaryotic and prokaryotic promoter elements. These sophisticated models excelled in generating a lot of functional promoters by assimilating and analyzing the information of distribution patterns inherent in extensive native promoter sequences (19,20). For example, synthetic promoters were *de novo* generated by Generative Adversarial Networks (GANs) and 70.8% generated promoters showed activities in *E. coli*, utilizing a 14,098 experimentally validated promoters as a training dataset (21). In a recent advancement, this model has been refined to not only produce entire promoter sequences but also to generate flanking regions while maintaining the integrity of the core regions in *E. coli* and mammalian cells (22). Meanwhile, a Variational Autoencoder (VAE) was trained to generate synthetic promoters of varying efficiency in cyanobacteria using 3,712 promoter sequences that identified through differential RNA sequencing (dRNA-seq) (23). These models demonstrated that robust performance could be achieved in the presence of extensive and high-quality experimental data. As the scope of metabolic engineering broadens to encompass an array of non-model prokaryotic organisms, the dearth of comprehensive and high-quality datasets emerges as a formidable challenge.

Recently, the field of text generation has witnessed significant advancements through the application of transfer learning, which combined Generative Pre-trained Transformers (GPT) with instruction datasets for the purpose of fine-tuning (24–26). This approach was exemplified by the success of ChatGPT (**Figure 1A**). In the pre-training phase, the model learns general and distribution patterns from a multitude of unlabeled datasets. Thereafter, a fine-tuning process, steered by data related to specific tasks, equips the model to efficiently adapt to a variety of application contexts (26). This methodology was particularly efficient in scenarios where data was limited, as transfer learning effectively augmented the precision of model and adaptability in such data-scarce situations (25). Biological sequences including proteins and nucleotides are regarded similar to natural languages as both of them use a set of symbols to represent information (27,28). Some nucleotide language models, e.g. DNABert, Nucleotide Transformers, GENA-LM, HyenaDNA (27–29), were designed to understand the genetic information. Transferrable understandings of sequences from genome could be learned by nucleotide language models, which were trained on existing genome sequences. However, most nucleotide language models are BERT-like models trained to learn representations of nucleotide sequences rather than to generate new sequences (30). One of the current challenges in developing generative model is to benchmark the quality of the generated sequences. Typically, the evaluation of sequence quality relies on scores of third-party *in silico* models and experimentally verification *in vivo* or *in vitro*. *In silico* evaluations commonly have a much higher throughput than experiments *in vivo* or *in vitro*, but experimental data are more reliable (31).

**Figure 1.**
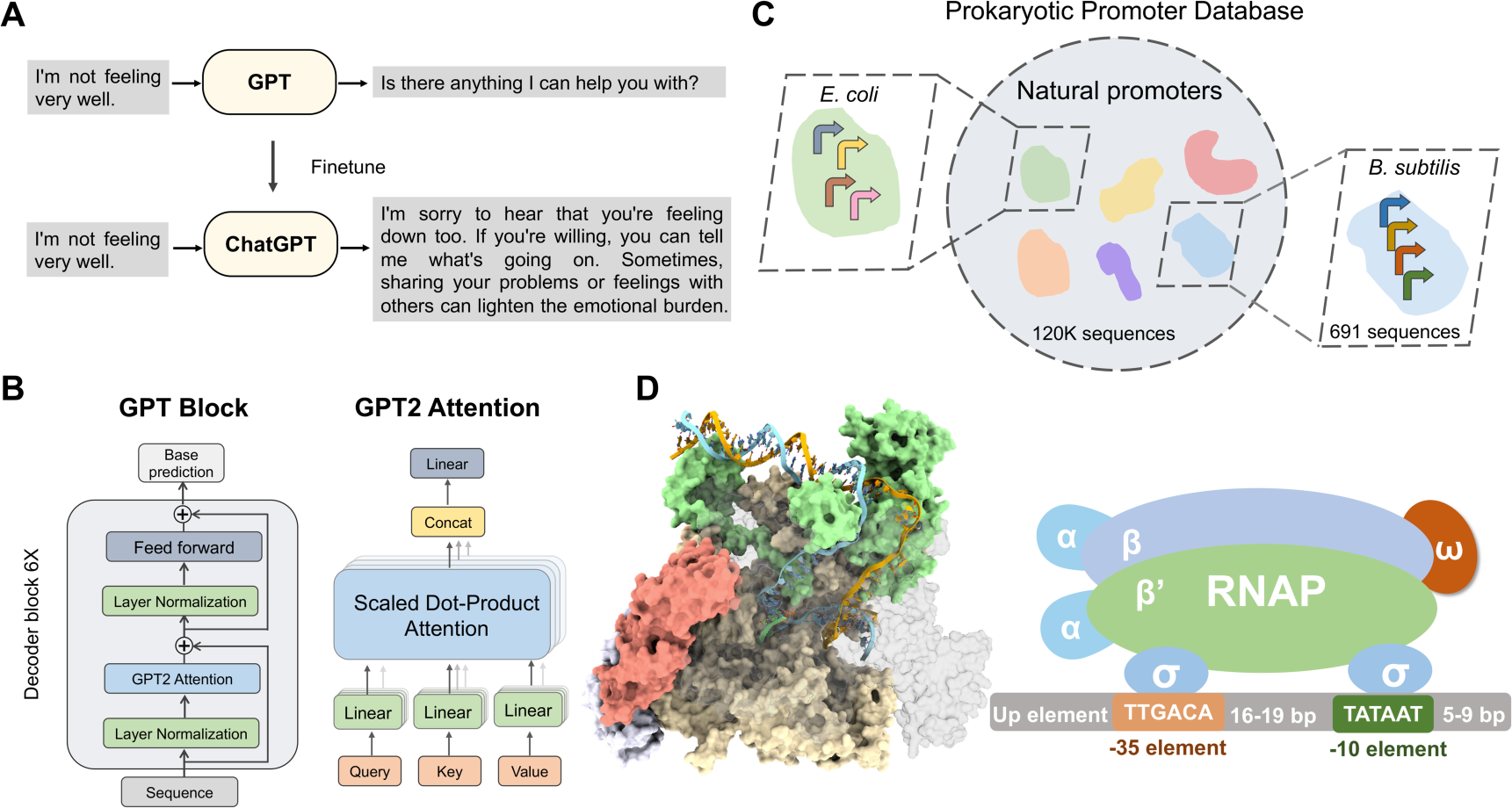
The framework of synthetic promoter design based on nucleotide language model. (A) A pretrained GPT model is capable of generating accurate sentences but may not always align with specific intentions or contexts. By finetuning a pretrained GPT, it can be adapted to not only produce correct sentences but also ensure alignment with the desired intent or context-specific requirements. (B) PromoGen-pre is a 7.4 million parameter neural network based on Transformer architecture, which relies on the self-attention mechanism to capture the distribution of tokens conditioned on context. (C) The PromoGen-pre is trained to generate artificial nucleotide sequence by minimizing the cross-entropy loss on the PPD database. (D) A diagrammatic representation depicts the formation of a transcription initiation complex, comprising promoters and RNA polymerase (RNAP). Promoters are primarily composed of up element, −35 element, flanking sequences, and −10 element. During the assembly of the transcription initiation complex, the α subunit is responsible for anchoring RNAP to the promoter region. The β and β’ subunits are involved in recognizing the sigma factor, while the ω subunit assists in the assembly of RNA polymerase.

To address the challenge of insufficient promoter data of certain species, we designed a pre-trained model based on Transformer, termed the PromoGen-pre. This model was further refined with species-specific promoter data to yield 27 distinct promoter-generating models for both model and non-model species. To validate the effectiveness of our strategy, we evaluated the key characteristics of the generated promoters including position weight matrix, 6-mer frequency correlation and −10 region distribution in *E. coli* and *B. subtilis*. We developed a scoring model to evaluate the *E. coli* promoters generated by our model and WGAN-GP, demonstrating the competitiveness of our model. Most of the generated promoters exhibited confirmed activities in *B. subtilis*. An online webserver was developed to enhance the accessibility of the promoter-generative models where users could generate *de novo* promoters for 27 different species based on the PromoGen model. The related models and analysis methods were listed in Supplementary Table S1. Via aforementioned transfer learning approach for generating new promoters from scratch, we expanded the available promoters for a specific species and offered potential candidates for use in metabolic engineering even when the available data was limited.

## Results

### PromoGen-pre is a generative pre-trained model for promoter sequences

Unsupervised language models have become the foundation of current advancement in the natural language processing (NLP) field (32). Language model could be built and trained based on the large number of unlabeled data, and be improved via transfer-learning for performing downstream tasks with task-specific dataset. Transfer-learning dependent strategy, which has been successfully exploited to generate functional proteins across diverse families recently, has yet been reported to generate new functional DNA elements. To investigate the potential of transfer learning on generating DNA regulatory elements, we developed a decoder-only Transformer-based model, termed PromoGen-pre, to specifically capture the characteristics of prokaryotic promoter sequences (**Figure 1B**). Unlike the conventional byte-pair encoding (BPE) or k-mer tokenization strategy (33,34), the DNA sequences were tokenized at the character level in PromoGen-pre to ensure that the original location of each base in the DNA sequence was kept within the position of each token, allowing the DNA generation more controllable (**Supplementary Figure S1A**). The tokenized input sequence was embedded as numerical representation *X_emb_* ∈ ℝ^*L×D*^, where *L* is the sequence length and *D* is the dimension of the hidden vector of each token. PromoGen model further updated the numerical representation though GPT2Blocks with multi-head self-attention (**Figure 1B, Supplementary Figure S1B**) (25).

When predicting the next token, the long-distance relation across the sequence would be taken account by PromoGen-pre via the self-attention mechanism. PromoGen-pre was trained by autoregressive approach, where the probability of each nucleotide in the sequence was conditioned solely on the upstream nucleotides. Cross-entropy loss was utilized to evaluate and adjust the performance of PromoGen-pre during the model training process. The 10 million tokens from more than 129k sequences that containing experimentally verified promoters and Transcriptional Start Sites (TSS) in the Prokaryotic Promoter Database (PPD), which is one of the most comprehensive promoter sequence databases containing sequences from more than 27 species, were subjected to the pretraining of PromoGen **(Figure 1C, D)** to ensure the high quality and reduced deviation (**Figure 2A, 2B**) (35).

**Figure 2.**
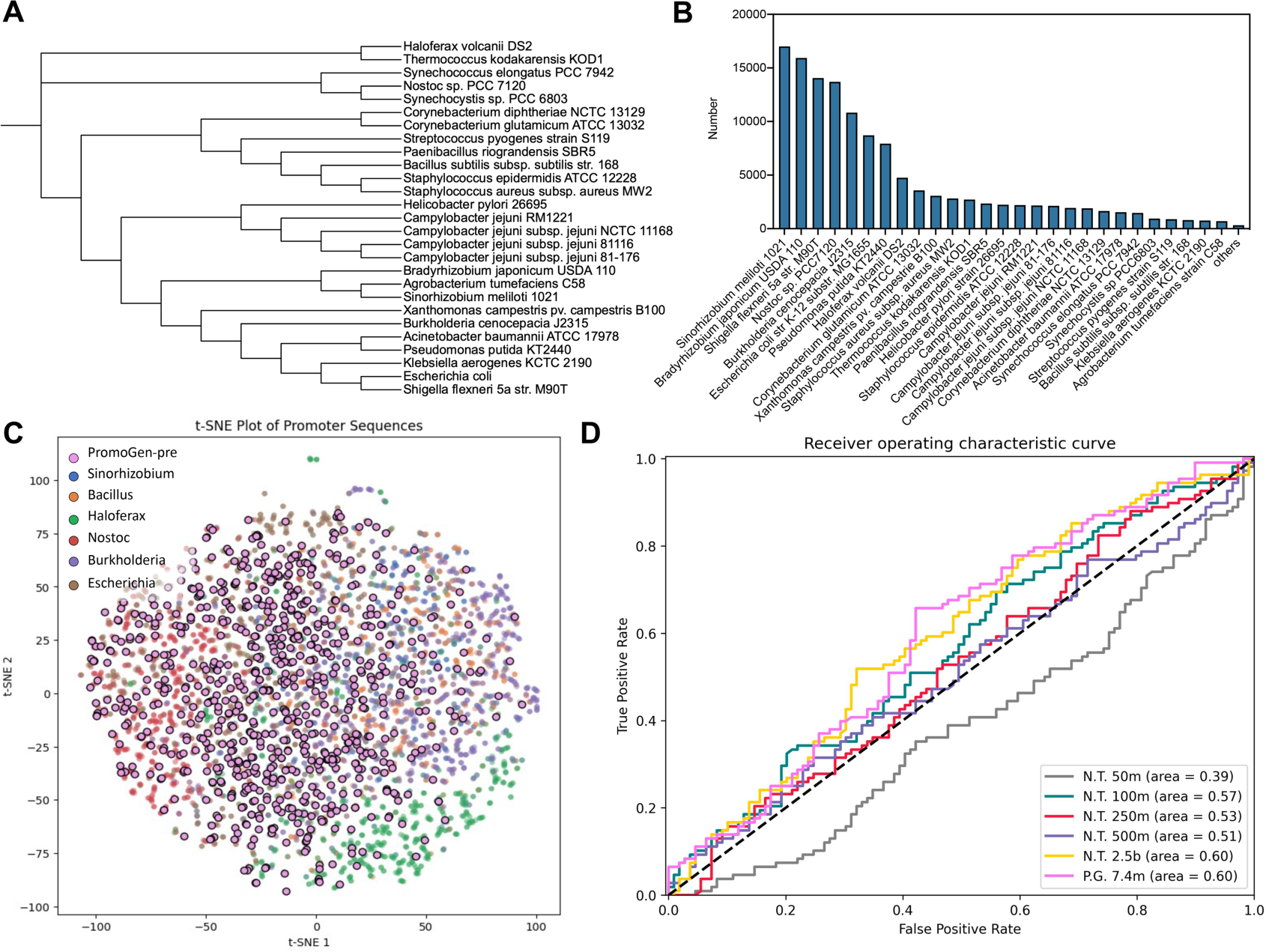
The training of PromoGen-pre. (A) Evolutionary tree of 27 major species included in PPD. (B)The number of promoter sequences contained in each species within PPD. The others indicate the sum of promoters from species except the 27 major ones. (C) The t-SNE, employed as a dimensionality reduction technique, was utilized to illustrate the spatial distribution of both native and generated promoters in a two-dimensional space. The distances between points represent the similarity. Blue, orange, green, red, purple, and grey color points represent the promoter sequences of *Sinorhizobium*, *Bacillus*, *Haloferax*, *Nostoc*, *Burkholderia*, and *Escherichia*, respectively. The pink color points represent the generated sequences of PromoGen. (D) The model likelihoods of PromoGen-pre can accurately predict the activity of generated promoters compared with different parameter sizes of pre-trained nucleotide transformers.

After the above training, we observed that the promoters from the same species were clustered close to each other when the sequence representations encoded by PromoGen-pre model were projected to a two-dimensional plan, indicating that the model has learned to distinguish the promoters by species in the absence of supervision related to species information (**Figure 2C**). For example, the sequences from *Haloferax* were notably well distinguished from other species by PromoGen, highlighting the differences of regulatory elements between Archaea and Bacteria (**Figure 2C**) and confirming that PromoGen-pre is able to encode meaningful promoter representations. On the other hand, some projected points representing promoters from different species were also gathered together in the Figure 2C, indicating that some promoters from distinguished species are sharing common patterns which is in agreement with the well-known observation that bacteria are able to recognize certain heterologous genetic elements. This observation reinforced our rationale to tackle the low-data challenge by transfer learning.

Since PromoGen-pre has been trained to obtain meaningful representations from native promoters, we investigated whether the trained PromoGen-pre could generate functional artificial promoters. The results showed that the projected points of artificial promoters generated by PromoGen-pre (pink points in Figure 2C) was dispersedly distributed within, and did not form obvious cluster outside of, the ones of native promoters, indicating that the generated promoters were likely to be functional as the native ones. So far, the generated promoters were not species specific although the pre-training stage allowed PromoGen-pre to learn the comprehensive features of promoters from different species, encompassing a broader spectrum and evolutionary information.

Although previous works developed a collection of benchmarks including eighteen genomic related prediction tasks (27), we find it is still challenging to evaluate whether nucleotide language models could understand prokaryotic promoters because currently only mouse and human promoters were involved in the benchmarks. On the other hand, only promoter sequences, but not genomic chunks, were provided to PromoGen-pre for training, making PromoGen-pre being intrinsically capable of distinguishing native promoters and random sampled genomic chunks, which was a challenging task for other models. Therefore, a benchmark dataset based on experimentally verified synthetic promoters was needed for PromoGen-pre.

We first evaluated the ability of PromoGen-pre to distinguish whether a generated sequence is a promoter in a zero-shot manner (36). The log-likelihood of each sequence was calculated by PromoGen-pre to a predicted score. As a comparison, we also evaluated the performance of the nucleotide transformer family models (NTFM) which have number of parameters range from 50 million (M) to 2.5 billion (B) NTFM enabled precise prediction of molecular phenotypes based on learned patterns from DNA sequences (27). We computed the area under the receiver operating characteristic (auROC) curves to predict the binary label from scores (37). The results showed that the PromoGen-pre, a small model with solely 7.4 M parameters, has log-likelihood auROC of 0.60, equally well, (if not better than) as the same level of the 2.5 B model (**Figure 2D**), better than the Nucleotide Transformer 50 M, 100 M, 250 M, and 500 M. This result showed that a relatively small amount of project-related high-quality data could fulfill the task, enabling PromoGen to reach the performance of large model trained with large dataset.

### Finetune the pre-trained model based on datasets from 27 species

PromoGen-pre has been trained on multiple prokaryotic promoter sequence datasets, endowing it with the capability to generate promoters for various species. To adapt the model for species-specific applications, we conducted fine-tuning based on the datasets of 27 major species in the dataset, resulting in the development of 27 distinct promoter models, encompassing species such as *E. coli*, *B. subtilis*, *Sinorhizobium*, *Corynebacterium*, and so on. Such transfer learning methodology facilitated the development of a robust promoter generative model even with relatively small-size datasets.

Taking *E. coli* model as an example to evaluate the effectiveness of transfer learning, we then tried to score and compare the generation quality of PromoGen-eco and a GAN-based WGAN-GP model through independent predictive models which were designed to predict activities of promoters. To select the most reliable predictive model for predicting the activities of the generated promoters, we evaluated twelve machine learning based predictive models by calculating metrics including MAE, RMSE, Spearman correlation coefficient, and Pearson correlation coefficient (**Supplementary Figure S2**). the Bayesian Bridge model was selected as our final predictive model because it consistently demonstrated exceptional performance no matter which calculation methods were used for evaluation. The Bayesian Bridge model achieved a Spearman correlation coefficient of 0.69 or 0.58, and a Pearson correlation coefficient of 0.70 or 0.60 on training and test data, respectively, demonstrating the accuracy of this model (**Figure 3A, 3B**).

**Figure 3.**
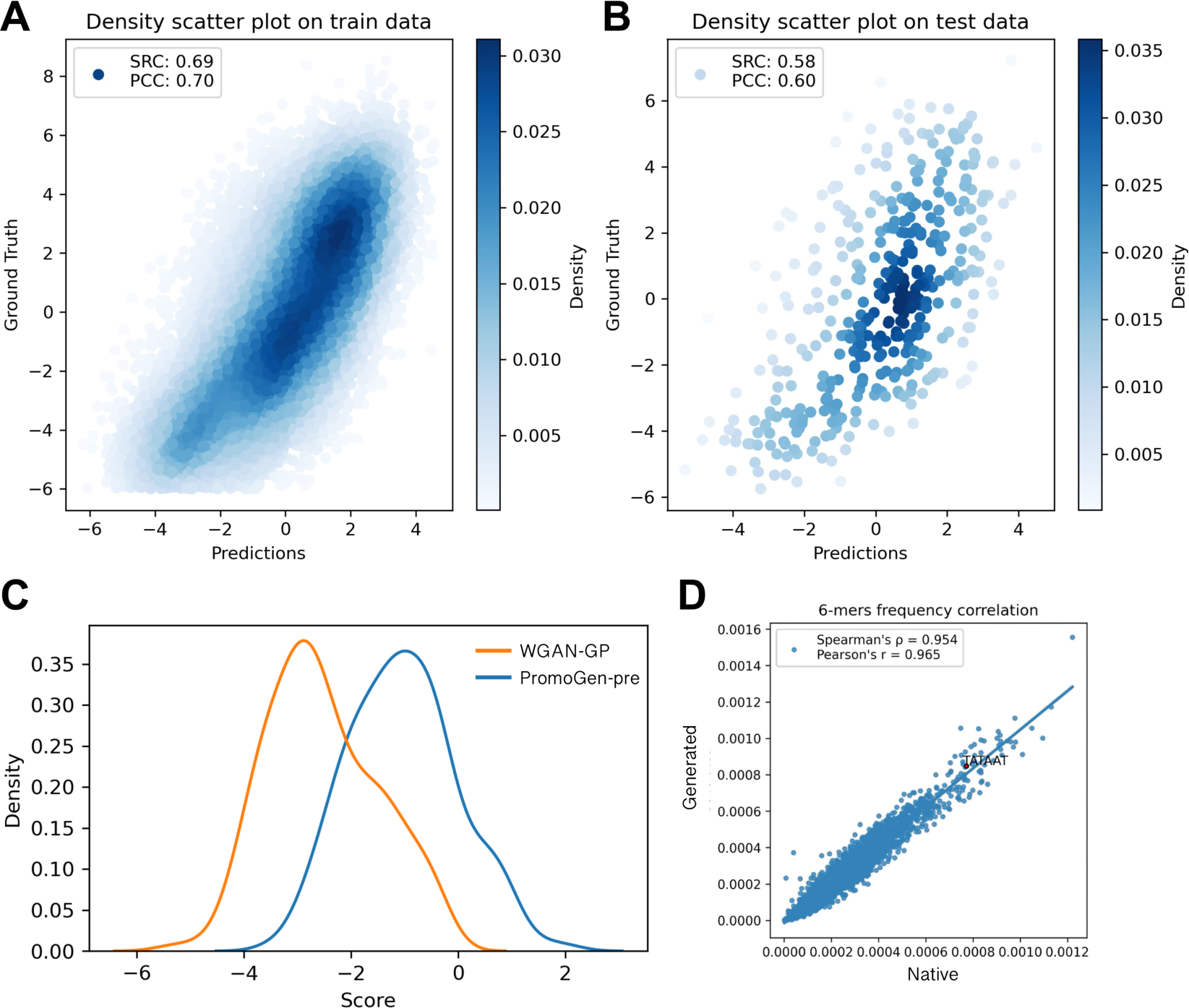
Comparison of PromoGen-eco with WGAN-GP model. (A) Density scatter plot on train data by Bayesian Bridge prediction model. The depth of color points represents density, with the x-axis and y-axis indicating predicted and actual promoter strengths, respectively. (B) Density scatter plot on test data by Bayesian Bridge prediction model. (C) The distribution of scores in PromoGen-eco and WGAN-GP. The orange and blue color linear represent WGAN-GP and PromoGen-eco, respectively. (D) The 6-mers frequency correlation between native and PromoGen-eco generated promoter sequences. Each data point represents a specific 6-mers, with the x-axis and y-axis indicating the frequency of the 6-mer in native and generated promoters, respectively.

We generated 200 promoter sequences by PromoGen-eco and WGAN-GP, individually, and used Bayesian Bridge prediction model to predict the promoter activities of these 400 promoters (21). As depicted in figure 3C, the average score of PromoGen-eco generated promoters was significantly higher compared to the ones generated by WGAN-GP, indicating the strong capability of PromoGen-eco in *de novo* promoter generation. In addition, another prediction model employed in previous study (21), Deep Promoter CNN, was utilized to assess the performance of PromoGen-eco. The results indicated that PromoGen-eco exhibited high competitiveness when compared with WGAN-GP (**Supplementary Figure S3**). Furthermore, analysis utilizing PWM revealed that the promoters generated by PromoGen-eco exhibited a distribution identical to that observed in native promoters while the distribution of promoters generated by WGAN-GP displayed slight variations from the native distribution (**Supplementary Figure S4**). To evaluate the similarity between natively occurring promoters and those synthesized by PromoGen-eco, we counted the hexamer frequencies of the promoters. The resulting Pearson and Spearman correlation coefficients were 0.954 and 0.965, respectively, indicating a high degree of similarity in the 6-mer frequency between the generated and native promoters (**Figure 3D**). These findings collectively suggested that PromoGen-eco has successfully captured the characteristics and distribution patterns of *E. coli* promoters, showcasing its strong capability in *de novo* promoter generation.

### Performance evaluation of PromoGen-bsu model

To further validate the effectiveness of various species specific PromoGen models, we performed promoter prediction and activity evaluation in *B. subtilis,* a species with higher promoter selectivity and is an important Gram-positive bacterium used in protein production and chemical manufacturing. We fine-tuned PromoGen-pre using 691 promoter sequences from *B. subtilis* to obtained PromoGen-bsu, which was then utilized to generate *B. subtilis* specific promoters (**Supplementary Figure S5**). To assess the quality of generated promoters, PromoGen-bsu generated promoters were randomly sampled and compared with native ones and PromoGen-pre generated ones. We utilized PWMs to depict the sequence motifs within the promoter regions derived from native sources, as well as those synthesized by PromoGen-pre and PromoGen-bsu (**Figure 4A**). Our investigation revealed a conserved motif within the −10 box, extending from position −13 to −8 in the native promoters. The conservation of this motif aligns with earlier studies conducted on a variety of bacterial species, including *B. subtilis*, *E. coli*, and cyanobacteria. Notably, in the −10 region of the PromoGen-bsu-generated promoters, we detected a sequence pattern mirroring that of the native promoters, suggesting that PromoGen-bsu has successfully replicated the essential features of this −10 motif. Additionally, the −35 region, located at the positions of −36 to −31, exhibited partial conservation in both the natively occurring and PromoGen-bsu-constructed promoter sequences. Contrasting with this, the sequence of PromoGen-pre attributes exhibited substantial deviation from the native *B. subtilis* promoters, exhibiting only a marginal preference for T, A bases within the −10 region and no preference within −35 region.

**Figure 4.**
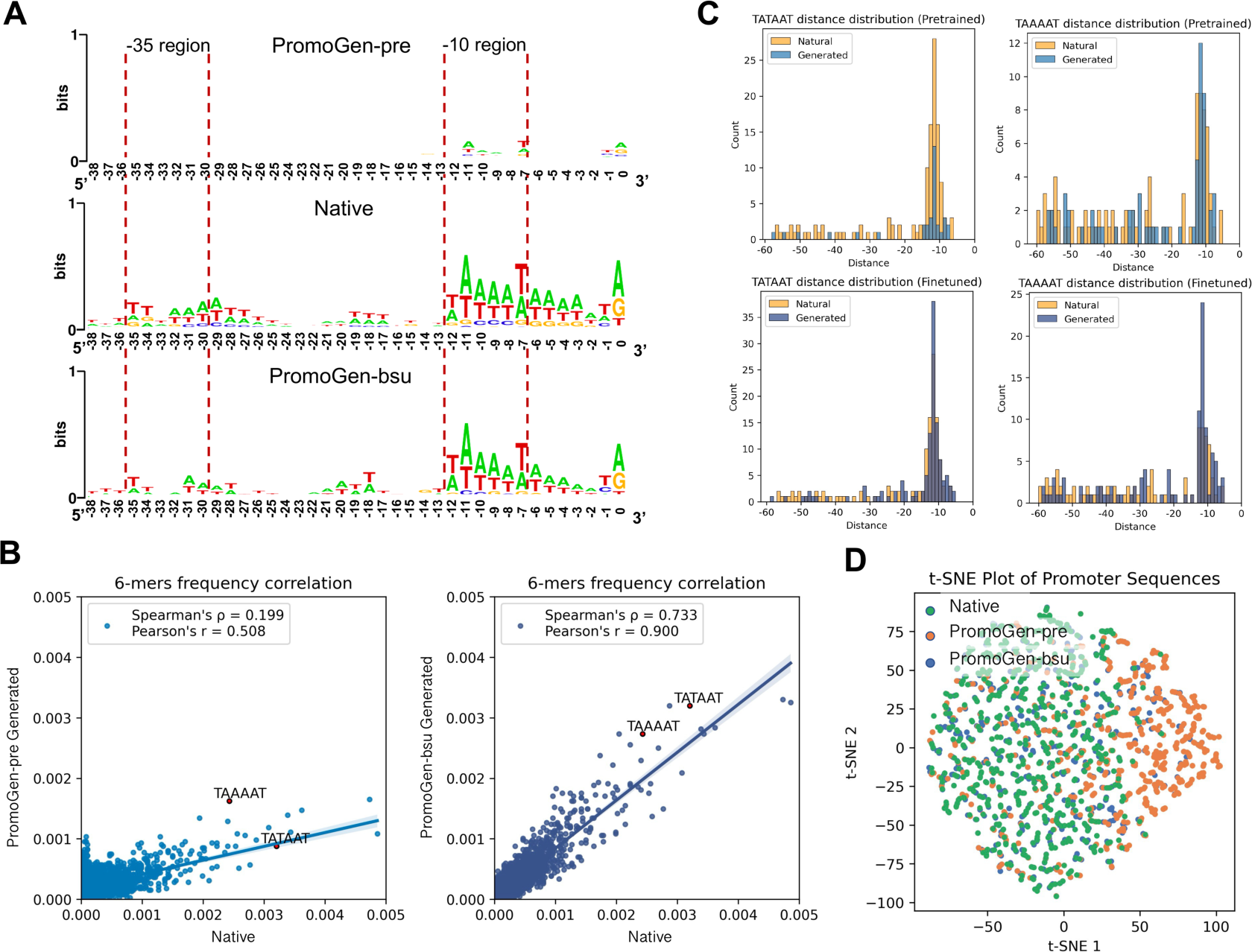
Evaluation of the PromoGen-pre and PromoGen-bsu generated promoters. (A) The sequence logos of the PromoGen-pre-generated, native, and PromoGen-bsu-generated promoters. The 0 represents the transcriptional start site (TSS). The key regions −10 and −35 have been marked by red dotted lines. (B) Spearman and Pearson values were used to assess the frequency of 6-mers between native and generated promoters. Each data point represents a specific 6-mers, with the x-axis and y-axis indicating the frequency of the 6-mer in native and generated promoters, respectively. (C) The distribution patterns of the top two 6-mers in PromoGen-pre and PromoGen-bsu within the −10 regions. The x-axis represents the distance from the TSS, and the y-axis represents the count of the 6-mer at that specific location. (D) The native and generated promoters are visualized in a two-dimensional space, with high-dimensional data condensed through t-SNE. The distances between points reflect the degree of relevance. Native promoters are depicted as green dots, while PromoGen-pre and PromoGen-bsu generated promoters are represented by orange and blue dots, respectively.

We conducted a quantitative assessment of hexamer frequencies in both natively occurring promoters and those generated by PromoGen-pre and PromoGen-Bsu. The analysis revealed a moderate correlation between the hexamer frequencies of native promoters and those generated by PromoGen-pre, as indicated by Pearson and Spearman correlation coefficients of 0.508 and 0.199, respectively. Conversely, a significantly higher correlation was observed in sequences produced by PromoGen-bsu, with Pearson and Spearman coefficients of 0.900 and 0.733, respectively, illustrating a closer alignment with the hexamer frequency patterns found in native promoters (**Figure 4B**). We noted a substantial consistency in high-frequency 6-mers between the native and PromoGen-bsu generated promoters, specifically including sequences like TAAAAT and TATAAT that encompass the −10 box. We also investigated the distance distribution of TAAAAT and TATAAT sequences relative to the transcription start site (TSS) (**Figure 4C**). The analysis of the promoters synthesized by both PromoGen-pre and PromoGen-bsu revealed a distinguishable accumulation of hexamers, particularly in the −10 region. This region exhibited a significantly higher frequency of these sequences compared to other areas, aligning with the distribution observed in native promoters. The sequences generated by PromoGen-bsu having a higher portion of both TATAAT and TAAAAT as the −10 region comparing to PromoGen-pre. These findings underscored the capability of the PromoGen-bsu model to accurately replicate the spatial distribution patterns of conserved motifs inherent to native promoters, thereby affirming its efficacy in reflecting the native positional preferences of these sequences.

The t-SNE visualization in Figure 4D illustrated the arrangement of promoter sequences, *i.e.* native, PromoGen-pre, and PromoGen-bsu, which were projected from a high-dimensional landscape into a two-dimensional plane. In this representation, the promoters generated by the pre-trained model, depicted as orange dots, were dispersed throughout the space, predominantly occupying the right side, which suggestsed a distinct pattern from the rest. In contrast, the native and PromoGen-pre -generated promoters, represented by green and blue dots respectively, were intermingled and uniformly situated on the left side. This indicated a shared distribution pattern among them, showing the PromoGen-bsu has learned the related distribution in *B. subtilis* promoters.

### The activity of generated promoters in *B. subtilis*

To assess the functionality of the synthesized promoters, we utilized PromoGen-bsu to create 50 *de novo* designed promoters, each has 61 nucleotides in length (**Figure 5A**). From this pool, 22 were arbitrarily chosen for experimental testing. As benchmarks, we employed native promoters from *B. subtilis*, such as P_43_, P_veg_, and P_lepA_, alongside three random sequences to serve as controls, both positive and negative. We constructed 28 shuttle vectors expressing fluorescent proteins *sfGFP*, each harboring the replication origins pBR322 and RepA for *E. coli* and *B. subtilis*, enabling *sfGFP* expression in both hosts. The activities of the promoters were determined by the fluorescence intensity emitted by *sfGFP*. Sampling was conducted during the logarithmic growth phase for fluorescence measurement. A fluorescence reading higher or lower than that of the randomly generated sequences indicated an active or inactive promoter, respectively.

**Figure 5.**
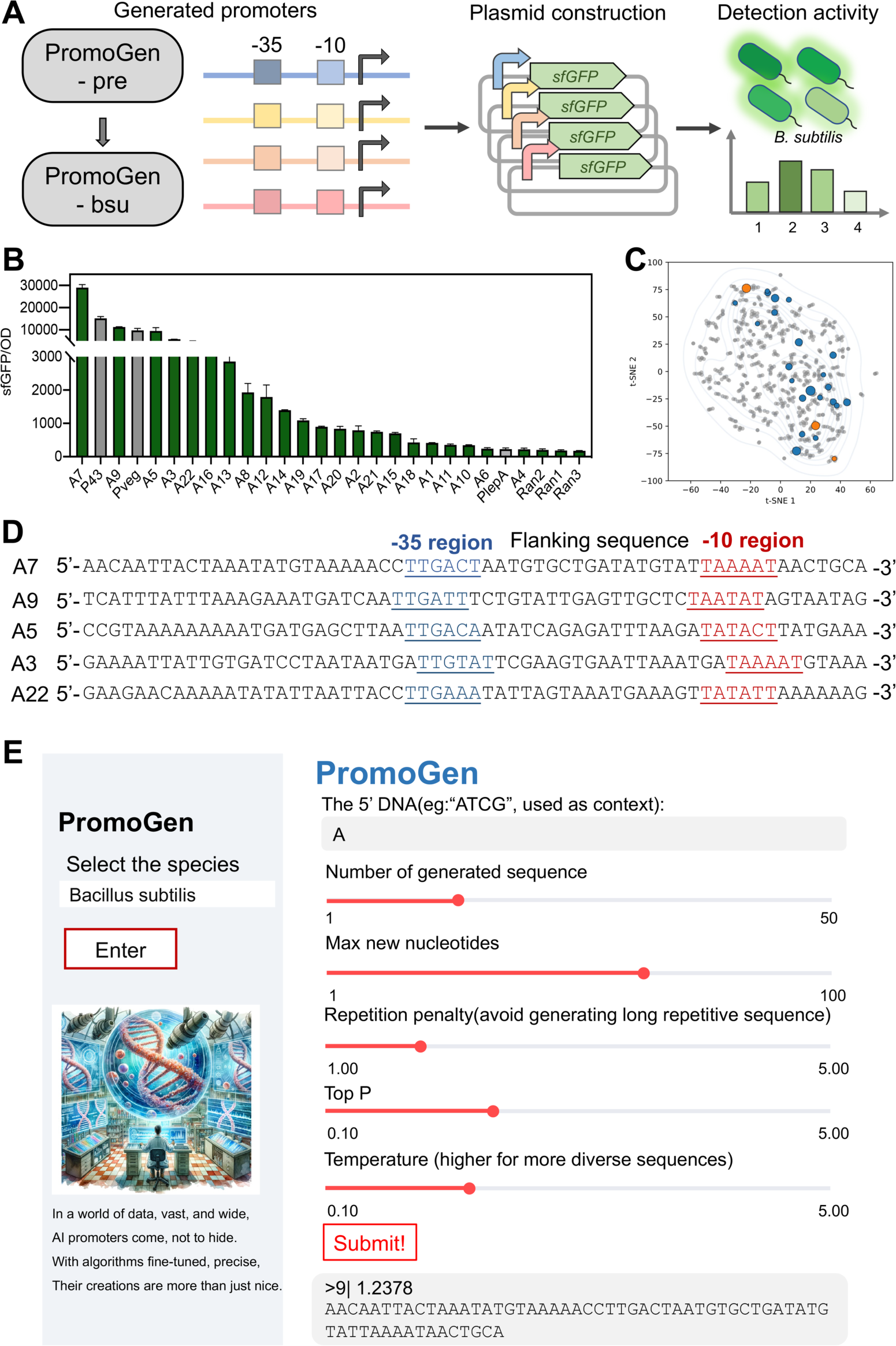
Validation of the generated promoter strength by *sfGFP* expression in *B. subtilis*. (A) Schematic representation of *sfGFP* expression and detection. (B) Strength of promoters were assessed by measuring the fluorescence intensity of *sfGFP* in *B. subtilis*. Twenty-two PromoGen-bsu generated promoters were detected, alongside three native promoters (P_43_, P_veg_, P_lepA_) serving as positive controls, and three dummy sequences as negative controls. (C) The native and validation promoters in the embedding spaces. The grey dots represent the 691 native promoters in the dataset. The orange and blue dots represent the native and generated validation promoters. The size of the dots shows the activity of the promoter. (D) The five most active promoters in the experiment were listed. Red and blue characters represent the −10 and −35 regions, respectively. (E) The web server of generate promoter sequences in 27 prokaryotes. When selecting the typical species, certain parameters could be adjusted, including the quantity and length of generated promoter sequences, base repetition penalty, top P and temperature.

As shown in Figure 5B, the results indicated that PromoGen-bsu could generate active promoters although the strength of the promoters vary. Three random sequences exhibited no activity. This result was expected because the range of promoter intensities use in the training data vary also. Almost all of these promoters are active, with about 18% of them including A7, A9, A5, and A3 showed activities similar to the natively strong promoters P_43_ and P_veg_. Notably, the A7 promoter stands out, exhibiting an activity rate 91% higher than that of P_43_. The strength of the other promoters varies, allowing for different levels of gene expression to be achieved. The bacterial strains expressed the *sfGFP* gene were subjected to observation under a fluorescence microscope. Notably, these organisms demonstrated notable fluorescence when compared to counterparts utilizing random sequences as promoters (**Supplementary Figure S6**). Furthermore, our investigation encompassed the analysis of t-SNE distribution patterns, compared experimentally tested 22 promoters with 691 natively promoters in *B. subtilis*. This analysis showed that the generated promoters were quite dispersed and were significantly distant from strong promoters, indicating a considerable difference (**Figure 5C**). Examination of the five most potent promoters uncovered a shared sequence, TTG, within the −35 region, and adjacent sequences were identified spanning positions 17-18 bp between the −35 and −10 regions (**Figure 5D**). Additionally, the activities of these promoters were tested in *E. coli.* The experimental results demonstrated that all promoters were active in *E. coli*, with 64% showing enhanced activity compared to *B. subtilis* (**Supplemental Figure S7**). This finding further validates that our model generates sequences that are not random but functionally significant in biological contexts. Moreover, these results support the concept of the universality of prokaryotic promoters, as discussed in studies from 2018. Our research also indicates that promoters in *B. subtilis* exhibit greater selectivity compared to those in other species.

To assess the similarity of the generated promoters to native promoters, a comprehensive BLAST analysis was performed, comparing the experimentally validated promoters against the entire genome of *B. subtilis*. The results showed that the generated sequences do not have similarity. This finding implies that the sequences were predominantly *de novo* designed by the models, rather than being a result of data leaks. This suggested that PromoGen-bsu has effectively captured the distribution patterns of active *B. subtilis* promoters, exploring a sequence space of 4^61^ and showcasing the ability to generate new active species-specific promoters.

### Development of a user-friendly web sever

To facilitate the rapid generation of promoter sequences, we have developed an online web server (https://promogen1.cloudmol.org/) providing fine-tuned models for 27 prokaryotic organisms. As shown in **Fig. 5E**, the promoter sequences could be generated after choosing the specific organism. The number and length of the generated promoters could be flexibly customized according to the needs. The generated results were presented in a sequence that ranked them according to decreasing values of the log-likelihood. The repetition penalty, Top P, and temperature were commonly utilized hyperparameters for controlling the diversity and quality of text generation. By increasing the repetition penalty, we could prevent the generative model from producing multiple repeated tokens. This strategy not only mitigates the occurrence of hallucinations in the generative model but also reduces the complexity involved in constructing plasmids. By adjusting the temperature and Top P settings, the diversity of the generated sequences could be regulated, thereby mitigating the risk of converging on local minima. This online platform offers a user-friendly interface for creating customized promoters for diverse species, serving specific needs and thereby facilitating the efficiency of subsequent experimental and research activities.

## Discussion

Regulatory elements play a crucial role in metabolic engineering (38). By selecting appropriate promoters, we could precisely control the gene expression of specific metabolic pathways, enhancing or inhibiting the fluxes of particular metabolic routes to increase the production of target chemicals. Identifying and verifying appropriate regulatory elements in understudied microbes is frequently a challenging and time-consuming task (18). Deep learning offers a promising approach for generating promoters from scratch or predicting the activity of existing promoters. However, the limited data available for understudied microbes poses a significant barrier to the effective utilization of deep learning models. Here, we proposed a fast and convenient solution based on generative self-supervised nucleotide language model, capable of generating promoters with different strengths. We employed a transfer learning strategy, which pre-trained the model using promoter sequences mainly from 27 species to learn the general patterns and paradigms of the promoters. Subsequently, fine-tuning was performed using species-specific promoter data, resulting in promoter models applicable to 27 different species. Our PromoGen was pretrained on multi-species data, making its adaptation to other species easier than WGAN-GP, which was only trained to generate *E. coli* promoters. Even for *E. coli* promoters, our PromoGen worked as accurate as, if not better, than the existing WGAN-GP model. Furthermore, under core region analysis and experimental validation, PromoGen-bsu model exhibited exceptional performance, with all of the generated promoters showing activities. Consistent results from model evaluation, key region analysis, and experimental validation indicated that PromoGen successfully learned promoter distribution patterns and could be used for generating entirely new promoters. For user convenience, an online webserver was developed containing data for 27 species, allowing users to quickly generate promoter sequences for their desired species.

The quality of the dataset plays a crucial role in the performance of deep learning models. Transfer learning offers an excellent solution for small-scale datasets. An increasing number of microorganisms are being used for various applications, but research on regulatory elements lags behind in non-model microorganisms. By utilizing the pre-trained model PromoGen-pre, we are able to reduce the data requirements for deep learning models, making it possible to *de novo* generate promoters for species with limited data. Although there are some differences in promoters among different species, the characteristics of the core region of prokaryotic promoters have been well-learned by the pretrained model PromoGen-pre. The activity of generated promoters could be enhanced by fixing key positions. Although the training data is limited comparing with other models, the information about promoter sequences is highly enriched. We observed DNA sequences differently from natural language in these aspects because there is no evidence that there is a pre-organized grammar across all species beside the Watson-Crick pairs. We also demonstrated that the zero-shot prediction accuracy of PromoGen-pre on promoter activities were comparable or higher than the ones of other nucleotide transformer models which are 6 to 330 times bigger than PromoGen-pre. We anticipated that this information enrich strategy would be helpful to projects with limited resources.

Although previous reports and our work demonstrated that it is effective to design promoters using generative models, it is challenging to assess the quality of generated promoters. The current methods for assessing generated sequences primarily encompass wet lab verification and computational experiment evaluation. Wet lab verification is the most precise approach, though large-scale experimental validations are time-consuming and labor-intensive. The evaluation methods for computational experiments include using comparative sample distributions and benchmark testing. Deep learning is capable of learning shallow features of data, which facilitates easy overfitting to match the sample distribution. Therefore, biologists should focus on establishing high-quality assessment datasets that are comprehensive, covering a wide array of scenarios and conditions, to guarantee the accuracy of evaluations.

Despite the fact that employing promoters of varying strengths is one of the most crucial methods for regulating gene expressions, the integration of diverse regulatory elements such as Ribosome Binding Sites (RBS), terminators, insulators, and others, offers expanded options for gene expression modulation (39–41). Furthermore, synthetic promoters, which establish a predetermined expression intensity, present challenges in dynamically regulating gene expression during growth. Promoters that are inducible or dependent on sigma factors could dictate the timing and intensity of gene activation. In the future, the design and development of pertinent models carry substantial promise for applications within the realm of synthetic biology.

In summary, we implemented a transfer learning strategy to address the challenges posed by a limited dataset. By the combination of pre-training and finetune, 27 species-generated models were developed for *de novo* design of promoter sequences, which were evaluated by prediction model and experiments validation as promoters with exceptional performance. Furthermore, we have developed a user-friendly web server that enables rapid generation of promoters. Our work provided a novel strategy and a universal pre-trained model PromoGen-pre, showcasing the potential of deep learning methods to explore the sequence space of regulation elements, especially in situations with limited data availability.

## METHODS

### Details of nucleotide language models

The vocabulary of PromoGen contains 7 tokens, including four bases A, T, C, G, and special tokens including [PAD], [MASK], and [UNK]. By applying one-hot encoding, the input sequences are transformed into numerical vectors that can be processed by neural networks. The one-hot vectors are transformed into hidden state vectors of 320 dimensions:

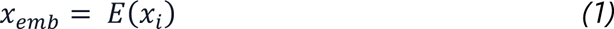

To distinguish the positions of each token, a learnable positional embedding is added:

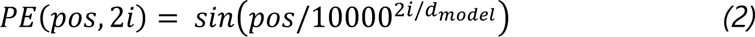

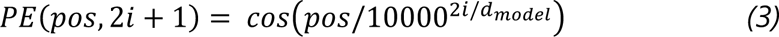

where the pos is the position of the context, i is the index of the dimension in the encoding vector, and the d_model_ represents the dimensionality of the outputs in all layers of the model. This position embedding is followed the original design of the GPT2 model (24).

The longest sequence allowed is 128. The hidden vectors are updated by 6 layers of GPT2 Block (25). In a GPT2 block, the hidden vectors are updated with multi-head self-attention, and a feed forward layer, with residual connections:

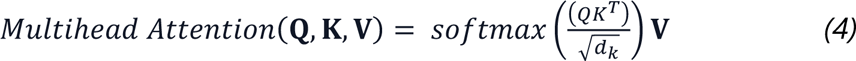

**Q** is query matrix, **K** is key matrix, **V** is value matrix, and *d_K_* is dimension of the key vectors. **W_1_** and **W_2_** is Weight matrices of the linear transformations. b_1_ and b_2_ is bias terms of the linear transformations. The number of head in multi-head self-attention is set to 4.

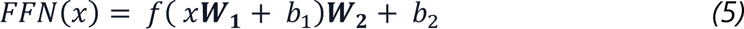

The feed forward layer (FFN) is consisted of two linear layers, to expend the dimension by a factor of 2 and activated using the Gaussian Error Linear Unit (GELU) function.

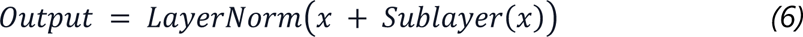

Residual connections and layer normalizations are applied in GPT2 block. The last hidden vectors are transformed to logits of tokens by the language modeling head, which is a linear layer has output dimension equal to the size of the vocabulary. The final output of the model is the probability of each base at the next position and generates promoter sequences by autoregression way from 5’ to 3’, one by one, which the next base has relied on all previously generated bases.

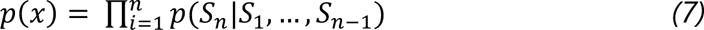

### Training of nucleotide language models

To train the PromoGen, we used 120k promoter sequences containing more than 27 species from the Prokaryotic Promoter Database (PPD). The PromoGen model is a model trained using the GPT-2 method (25). To efficiently train this model, the batch size was configured at 64, allowing for the simultaneous processing of 64 data samples. We predict the probability of each token based on previous context, comparing these predictions with actual data by computing cross-entropy loss to assess the prediction accuracy of model. To optimize the performance of model, the Adam optimizer was employed (42), renowned for its capability to dynamically adjust the learning rate, thereby bolstering the training efficiency and overall efficacy. We set several key parameters, including two decay rate parameters, β1 and β2, set to 0.9 and 0.999, respectively, and a constant called epsilon, set to 5e^-4^. During the fine-tuning phase, the weights from the pre-trained model are utilized as initial values. We configure the batch size to be 32, and the learning rate for the Adam optimizer is set at 1e^-5^. Other parameters remain consistent with those established in the PromoGen configuration.

### Training and selection of predictive models

We systematically evaluated 12 machine learning models to accurately predict the activity of promoters, including Extra Tree Regressor, Linear Regression, Gradient Boosting Regressor, Ada Boost Regressor, Ridge Lasso K-Neighbors Regressor, Random Forest Regressor, Bayesian Ridge, Support Vector Regression (SVR), eXtreme Gradient Boosting (XGB) Regressor, and Decision Tree Regressor (43–50). We extracted features from promoters using a nucleotide pre-trained model, employing the features from the [CLS] token as the representation for the entire sequence, which served as input to the machine learning models. To ensure a fair comparison of these methods, we conducted 5-fold cross-validation across the entire dataset ensuring independent predictions for all samples. At the same time, we calculated Mean Absolute Error (MAE), Root Mean Square Error (RMSE), Spearman correlation, and Pearson correlation coefficients (51,52).

To further enhance the accuracy of the predictive model, we conducted a hyperparameter search for the Bayesian Bridge model. These hyperparameters included maximum optimization iterations, alpha1, alpha2, lambda1, and lambda2, resulting in a total of 3^5^ combinations. Setting the maximum optimization iterations to 300 and alpha1, alpha2, lambda1 as 1e^-7^, and lambda2 as 1e^-5^ yielded the best performance in cross-validation. These optimal hyperparameters were used to Bayesian Bridge model for predicting promoter strength, ensuring our model achieves the highest level of accuracy.

### The model evaluation by position weight matrix (PWM) and 6-mer frequency analyses

We performed multiple sequence alignment separately for both native and model-generated promoter sequences. We calculated the weight for each position using a formula:

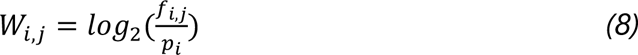

where *W_i,j_* represents the weight score of the *i*th base at position *j*. *f_i,j_* indicates the relative frequency of the base at position *j*, and *p_i_* represents the background frequency of that base. We utilized the obtained weight scores to create visualization using the online website WebLogo.

In the analysis of 6-mer frequencies, we conducted a comprehensive scan of all 6-mer sequences within both natural and synthetic promoters generated by PromoGen and PromoGen-bsu, subsequently computing the frequency of each individual 6-mer. During this process, particular attention was devoted to two pivotal 6-mer sequences, TATAAT and TAAAAT, which frequently play crucial roles in the functionality of promoters. To gain deeper insights into the representation of these key sequences across various promoters, we specifically calculated their abundance distribution compared with natural promoters.

### Plasmid construction

This investigation employed a variety of plasmids, detailed in Supplementary Table S2. All primer sequences were synthesized from Genewiz, located in Tianjin, China (Supplementary Table S3). The shuttle plasmid, pXY43-sfGFP, was utilized as a template, incorporating both pBR322 ori and repA ori for compatibility with *E. coli* and *B. subtilis* respectively. A series of 22 generated promoter sequences were integrated upstream of the superfolder green fluorescent protein (*sfGFP*) gene, replacing the existing P_43_ promoter via polymerase chain reaction (PCR) techniques. The amplified PCR constructs were subsequently cultured on LB agar plates containing 50 μg ml^−1^ of kanamycin to facilitate clone selection. Three clones were selected to Sanger sequencing for sequence verification in each plate. Furthermore, corresponding plasmids containing the P_veg_ and P_lepA_ promoters from *B. subtilis* were also engineered, alongside a set of plasmids with random sequences as promoters, to serve as negative controls in the study.

### Fluorescence measurements

The activity of these generated promoters was assessed based on the fluorescence intensity emitted by *sfGFP*. The relevant promoter-plasmid were transformed into *B. subtilis* strains, with single colonies subsequently cultivated in LB medium supplemented with 20 μg ml^−1^ kanamycin, maintained at 37°C and agitated at 220 r.p.m (Supplementary Table S4). After an overnight growth, these cultures were then transferred into fresh LB medium containing a 1% (v/v) inoculum, followed by an 8-hour incubation at 37°C. The fluorescence intensity was quantitatively measured using a microplate reader (BioTek) set at an excitation wavelength of 488 nm and an emission wavelength of 507 nm. Concurrently, the cell optical density (OD) was detected at 600 nm. All these experimental procedures were replicated thrice to ensure reliability.

## Data availability

All necessary data to assess the conclusions in the paper are presented within the paper itself and/or the Supplementary Materials.

## Author contributions

Y.-X.H., S.Y.G., and Y.X. conceptualized the project. Y.X. designed, built, and implemented deep models in this study. Y.X., and X.W.D. carried out the wet experiments. Y.-X.H., S.Y.G., and Y.X. analyzed the experiments data, and wrote the manuscript.

## Acknowledgements

This work was supported by the National Key R&D Program of China (2021YFC2100500), and the Natural Science Foundation of China (grant number 32370095). Part of the experiments was carried out in the Biological & Medical Engineering Core Facilities of the Beijing Institute of Technology. We would like to thank Jinyuan Sun for insightful discussions.

## Competing interests

The authors declare no competing interests.

## TOC

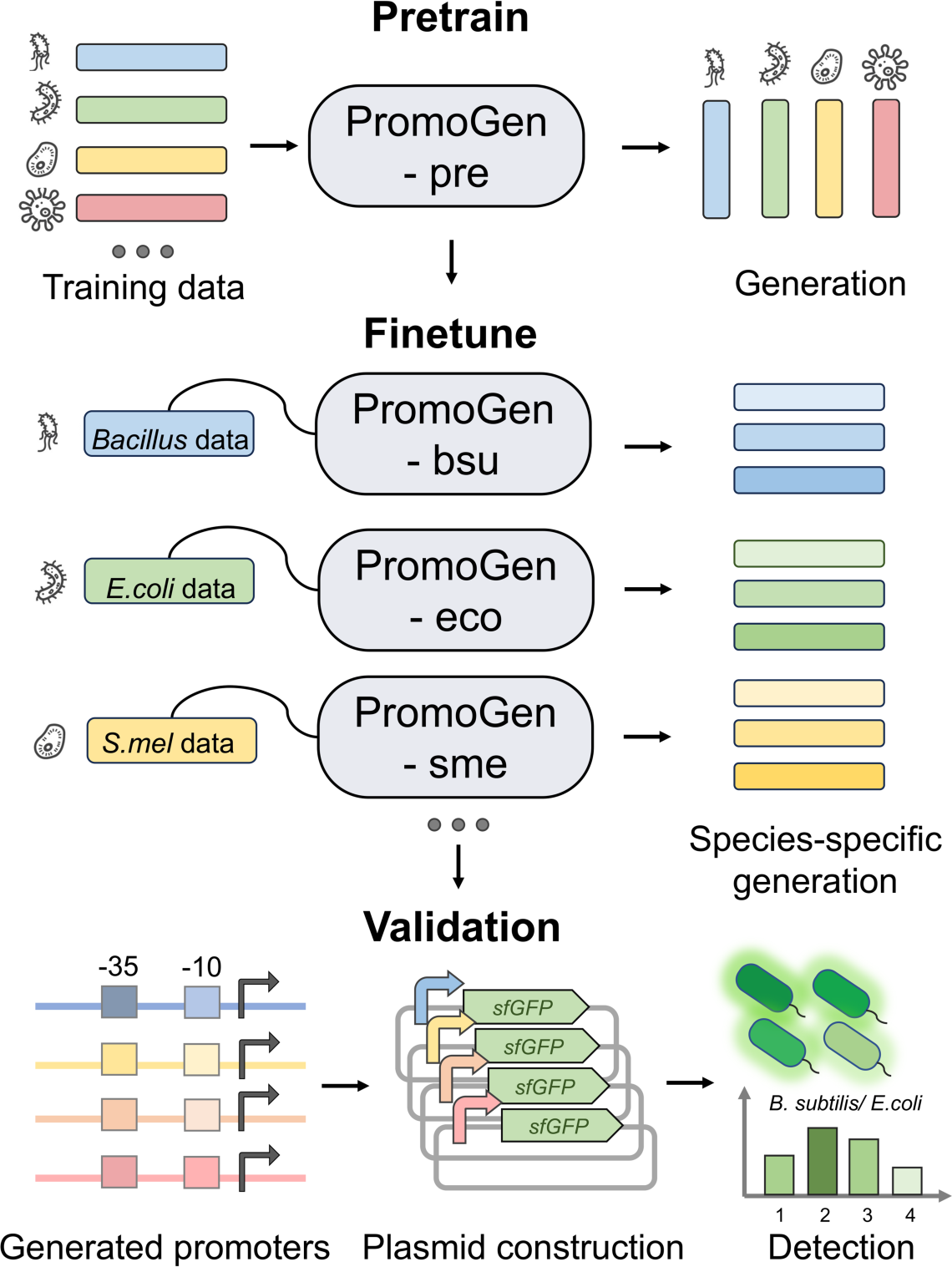

## Notes

### Competing Interest Statement

The authors have declared no competing interest.

### Summary of Updates

We have updated the introduction to make it smoother and more complete.

