## Supplemental information for "Species-specific design of artificial promoters by transfer-learning based generative deep-learning model"

**Table S1. The explanation of models and analysis methods.**

| Models/ methods | | Explanations |
|---|---|---|
| Generation models | PromoGen | A species-distinguishable nucleotide language model to generate species-specific functional promoters. |
| | PromoGen-pre | A pre-trained nucleotide language model based on Transformer. |
| | PromoGen-eco | Our nucleotide language model to generate *E. coli* promoters. |
| | PromoGen-bsu | Our nucleotide language model to generate *B. subtilis* promoters. |
| Tokenizer | Byte-Pair Encoding (BPE) | An algorithm primarily used in native language processing and text compression (1). |
| | k-mer tokenization | A method for breaking down these long biological sequences into smaller, manageable units for various types of analysis (2). |
| Prediction models | Bayesian Bridge regression | An advanced statistical technique that combines the concepts of Bayesian inference and Bridge regression and provides a probabilistic approach to regression analysis (3). The model was used to predict the promoter activity in PromoGen-eco and WGAN-GP (4). |
| | Zero-shot manner | A model is required to recognize or handle tasks, categories, or entities that it has not seen during training (5). It was used to score the 217 *B. subtilis* promoters. |
| Analysis methods | auROC | This is the area under the ROC curve. An auROC of 1 represents a perfect classifier, and an auROC of 0.5 represents a worthless classifier. The higher the auROC, the better the classifier is at distinguishing between the positive and negative classes (6). |
| Analysis methods | MAE | MAE measures the average magnitude of errors in a set of predictions and calculated as the average of the absolute differences between the predicted values and the actual values (7). |

| | | |
|---|---|---|
| Analysis methods | RMSE | RMSE evaluates how accurately a model predicts the target variable and is calculated by taking the square root of the MSE (7). |
| | Spearman | Spearman's correlation coefficient measures the strength and direction of association between two ranked variables (8). |
| | Pearson | Pearson's correlation coefficient is a key statistical tool for measuring linear relationships between two continuous variables (8). |

**Table S2. The plasmids used in this study**

| Plasmids | Description | Source |
|---|---|---|
| pXY-P43-sfGFP | A plasmid expressing sfGFP under the control of the P43 promoter | Laboratory stock |
| pXY-PlepA-sfGFP | A plasmid expressing sfGFP under the control of the PlepA promoter | This study |
| pXY-Pveg-sfGFP | A plasmid expressing sfGFP under the control of the Pveg promoter | This study |
| pXY-A1-sfGFP | A plasmid expressing sfGFP under the control of the A1 promoter | This study |
| pXY-A2-sfGFP | A plasmid expressing sfGFP under the control of the A2 promoter | This study |
| pXY-A3-sfGFP | A plasmid expressing sfGFP under the control of the A3 promoter | This study |
| pXY-A4-sfGFP | A plasmid expressing sfGFP under the control of the A4 promoter | This study |
| pXY-A5-sfGFP | A plasmid expressing sfGFP under the control of the A5 promoter | This study |
| pXY-A6-sfGFP | A plasmid expressing sfGFP under the control of the A6 promoter | This study |
| pXY-A7-sfGFP | A plasmid expressing sfGFP under the control of the A7 promoter | This study |

| | | |
|---|---|---|
| pXY-A8-sfGFP | A plasmid expressing sfGFP under the control of the A8 promoter | This study |
| pXY-A9-sfGFP | A plasmid expressing sfGFP under the control of the A9 promoter | This study |
| pXY-A10-sfGFP | A plasmid expressing sfGFP under the control of the A10 promoter | This study |
| pXY-A11-sfGFP | A plasmid expressing sfGFP under the control of the A11 promoter | This study |
| pXY-A12-sfGFP | A plasmid expressing sfGFP under the control of the A12 promoter | This study |
| pXY-A13-sfGFP | A plasmid expressing sfGFP under the control of the A13 promoter | This study |
| pXY-A14-sfGFP | A plasmid expressing sfGFP under the control of the A14 promoter | This study |
| pXY-A15-sfGFP | A plasmid expressing sfGFP under the control of the A15 promoter | This study |
| pXY-A16-sfGFP | A plasmid expressing sfGFP under the control of the A16 promoter | This study |
| pXY-A17-sfGFP | A plasmid expressing sfGFP under the control of the A17 promoter | This study |
| pXY-A18-sfGFP | A plasmid expressing sfGFP under the control of the A18 promoter | This study |
| pXY-A19-sfGFP | A plasmid expressing sfGFP under the control of the A19 promoter | This study |
| pXY-A20-sfGFP | A plasmid expressing sfGFP under the control of the A20 promoter | This study |
| pXY-A21-sfGFP | A plasmid expressing sfGFP under the control of the A21 promoter | This study |
| pXY-A22-sfGFP | A plasmid expressing sfGFP under the control of the A22 promoter | This study |
| pXY-ran1-sfGFP | A plasmid expressing sfGFP under the control of the ran1 promoter | This study |

| | | |
|---|---|---|
| pXY-ran2-sfGFP | A plasmid expressing sfGFP under the control of the ran2 promoter | This study |
| pXY-ran3-sfGFP | A plasmid expressing sfGFP under the control of the ran3 promoter | This study |
| pXY-J23100-sfGFP | A plasmid expressing sfGFP under the control of the J23100 promoter | This study |
| pXY- J23102-sfGFP | A plasmid expressing sfGFP under the control of the J23102 promoter | This study |
| pXY- J23119-sfGFP | A plasmid expressing sfGFP under the control of the J23119 promoter | This study |

**Table S3. The Primers used in this study.**

| Primers | Sequences (5’-3’) |
|---|---|
| A1-F | AGCTAATTATTTTATTATAATGATATATAATAAATATTAAAGGAGGTGATAAAAATGGTTAGC |
| A1-R | TCATTATAATAAAATAATTAGCTATTCAAAAATATTTTTGCTTTTGCTAGTTACCCTCGAGATCTGG |
| A2-F | CTTTGAATGAATATTTTACAATTGTTAATATATAAAGAAAAGGAGGTGATAAAAATGGTTAGC |
| A2-R | CAATTGTAAAATATTCATTCAAAGATTTTATAGAGCGTTTTTTGAACCTAGTTACCCTCGAGATCTGG |
| A3-F | GATTGTATTCGAAGTGAATTAAATGATAAAATGTAAAAAAGGAGGTGATAAAAATGGTTAGC |
| A3-R | AATTCACTTCGAATACAATCATTATTAGGATCACAATAATTTTCCTAGTTACCCTCGAGATCTGG |
| A4-F | GAAGTTTTTCATTACATATAAAAGCTATACTATACTCAAAAGGAGGTGATAAAAATGGTTAGC |
| A4-R | TTATATGTAATGAAAAACTTCAAGATTTGGTTAATTTTACCATTCTAGTTACCCTCGAGATCTGG |
| A5-F | AATTGACAATATCAGAGATTTAAGATATACTTATGAAAAAAGGAGGTGATAAAAATGGTTAGC |
| A5-R | AATCTCTGATATTGTCAATTAAGCTCATCATTTTTTTTTACGGCTAGTTACCCTCGAGATCTGG |
| A6-F | TTCGAAAGTTTATTAAATTCAATGTAATATAAATGATGAAAGGAGGTGATAAAAATGGTTAGC |
| A6-R | TGAATTTAATAAACTTTCGAAAATTTTACCATTGCAATTGCAACCTAGTTACCCTCGAGATCTGG |

| | |
|---|---|
| A7-F | ACCTTGACTAATGTGCTGATATGTATTAAAATAACTGCAAAAGGAGGTGATAAAAATGGTTAGC |
| A7-R | ATCAGCACATTAGTCAAGGTTTTTACATATTTAGTAATTGTTCTAGTTACCCTCGAGATCTGG |
| A8-F | TTATTCATAATGAGTGCAGTGCTATAATAAAAGTAAAAGGAGGTGATAAAAATGGTTAGC |
| A8-R | ACTGCACTCATTATGAATAATAATAATCAACACAATACGTAAAAAGCTAGTTACCCTCGAGATCTGG |
| A9-F | CAATTGATTTCTGTATTGAGTTGCTCTAATATAGTAATAGAAAGGAGGTGATAAAAATGGTTAGC |
| A9-R | ACTCAATACAGAAATCAATTGATCATTTCTTTAAATAAATGACTAGTTACCCTCGAGATCTGG |
| A10-F | TTGCATTGAGTTTTGCTCAAAAGGTGTAAAATAACAAAAAAAGGAGGTGATAAAAATGGTTAGC |
| A10-R | TTGAGCAAAACTCAATGCAAATATTTTCGTTATTTATTTATTCTAGTTACCCTCGAGATCTGG |
| A11-F | AACAATCATGCAAATTTTACTATAATACGATGAAATTTAAAGGAGGTGATAAAAATGGTTAGC |
| A11-R | GTAAAATTTGCATGATTGTTAAAATCATAATCAAAAACTTTTACTAGTTACCCTCGAGATCTGG |
| A12-F | TTTGTTATTGTGGTGAACATTGATATAATGACATAAAAGGAGGTGATAAAAATGGTTAGC |
| A12-R | ATGTTCACCACAATAACAAAAATAATGTTAATTTAATTAATTCAGACTAGTTACCCTCGAGATCTGG |
| A13-F | ATTGATTTAATTACACCTGACGTATTATATTTAATTTGAAAGGAGGTGATAAAAATGGTTAGC |
| A13-R | CGTCAGGTGTAATTAAATCAATGATATTTATAAATAATTATCAAGCTAGTTACCCTCGAGATCTGG |
| A14-F | TCTTGATATAAATCAAAGATATGCATAAAATGAATTTTGAAAGGAGGTGATAAAAATGGTTAGC |
| A14-R | TATCTTTGATTTATATCAAGAAAATATTCCATATCAAAATCATCTAGTTACCCTCGAGATCTGG |
| A15-F | AATTAACAAGAAGATTGAAAATTCTGATATAATGCATAAAAGGAGGTGATAAAAATGGTTAGC |
| A15-R | TTTTCAATCTTCTTGTTAATTATTTGGCTGTCATTTTTAGTTAACTAGTTACCCTCGAGATCTGG |
| A16-F | TAATTGTTAAATCGTTATTTTACGTTATAATAATAAAGAAAGGAGGTGATAAAAATGGTTAGC |
| A16-R | AAATAACGATTTAACAATTAAAGATTTCCATAGTAATGGAGTACTAGTTACCCTCGAGATCTGG |
| A17-F | ATGTTGACAATGAATTGTCACCGATGAATAATACAATAAAAAGGAGGTGATAAAAATGGTTAGC |
| A17-R | TGACAATTCATTGTCAACATACGTTCATTTTTTTTCATAAGGCTAGTTACCCTCGAGATCTGG |

| | |
|---|---|
| A18-F | CAAGTTGAAAACTTGCTAAAAGCTGATAAAGTAGAAGCTAAAAGGAGGTGATAAAAATGGTTAGC |
| A18-R | TTTTAGCAAGTTTTCAACTTGATTTTCAAAAATTTCACCCAACTAGTTACCCTCGAGATCTGG |
| A19-F | TGACTACATCATATTCAATCATAGTATAATAAAATTAAAAAAGGAGGTGATAAAAATGGTTAGC |
| A19-R | GATTGAATATGATGTAGTCAAATTTCACAAGTTTTAAGTATACTAGTTACCCTCGAGATCTGG |
| A20-F | AACATTGAAGTAAATTTGAAAAAGGCTATTATACAATAGAAAGGAGGTGATAAAAATGGTTAGC |
| A20-R | TTCAAATTTACTTCAATGTTTCAAATTTTTTTTGTTAAATACCTAGTTACCCTCGAGATCTGG |
| A21-F | GTAATTGACAAACACTGAAACAATGCATAATATATAAGCTAAAGGAGGTGATAAAAATGGTTAGC |
| A21-R | TTTCAGTGTTTGTCAATTACATAATTTTAGATAATTGTAGACTAGTTACCCTCGAGATCTGG |
| A22-F | TTACCTTGAAATATTAGTAAATGAAAGTTATATTAAAAAAGAAAGGAGGTGATAAAAATGGTTAGC |
| A22-R | ATTTACTAATATTTCAAGGTAATTAATATATTTTTGTTCTTCCTAGTTACCCTCGAGATCTGG |
| Test-1 | TACAGCACCTTCTAAAAGCGTTG |
| Test-2 | AAGCAGCAGATTACGCGC |
| lepA-F | TGTTTTACATTGAATCTTTACAATCCTATTGATATAATCTAAGCTAGTGTATTTAAAGGAGGTGATAAAAATGGTTAGC |
| lepA-R | TGTAAAGATTCAATGTAAAACAAGAAAGAGAAAAGTTCCCTATCATACTAGTTACCCTCGAGATCTGG |
| veg-F | AATTTTATTGACAACGTCTTATTAACGTTGATATAATATTGCAAGCTTGCAAAAAAGGAGGTGATAAAAATGGTTAGC |
| veg-R | TAAGACGTTGTCAATAAAATTATTTTGACAAAATTCTATGATTCCTCAACTAGTTACCCTCGAGATCTGG |
| Ran1-F | AGCTACGATGGGCTCCCGTACGTAGCTAGCTAGAGTTGCAAAGGAGGTGATAAAAATGGTTAGC |
| Ran1-R | TACGGGAGCCCATCGTAGCTAGTCACGTTTACTTTATCGACTCTAGTTACCCTCGAGATCTGG |
| Ran2-F | GGCTAGCTAGCCCGTGGGGACTAGCTAGCTACGCTAGGCAAAGGAGGTGATAAAAATGGTTAGC |
| Ran2-R | TCCCCACGGGCTAGCTAGCCTAGCTACGGGGAGCAAACTAGCCTAGTTACCCTCGAGATCTGG |

| | |
|---|---|
| Ran3-F | ATCCCCACTTTTTTCAGCTAGTACACGTGTGTGACTAGCTAAAGGAGGTGATAAAAATGGTTAGC |
| Ran3-R | CTAGCTGAAAAAAGTGGGGATCGATCAGCATTGTACGGTGGGCTAGTTACCCTCGAGATCTGG |
| J23100-F | GCTAGCTCAGTCCTAGGTATAATGCTAGCAAAGGAGGTGATAAAAATGGTTAGC |
| J23100-R | ATACCTAGGACTGAGCTAGCTGTCAAGAATTCTAGTTACCCTCGAGATCTGG |
| J23102-F | TAGCTCAGTCCTAGGTACAGTGCTAGAAAGGAGGTGATAAAAATGGTTAGC |
| J23102-R | CTGTACCTAGGACTGAGCTAGCCGTCAAGAATTCTAGTTACCCTCGAGATCTGG |
| J23119-F | CAGCTAGCTCAGTCCTAGGTACTGTGCTAGAAAGGAGGTGATAAAAATGGTTAGC |
| J23119-R | TACCTAGGACTGAGCTAGCTGTCAAGAATTCTAGTTACCCTCGAGATCTGG |

**Table S4. The strains used in this study.**

| **Strains** | **Description** | **Source** |
|---|---|---|
| *E. coli* JM109 | *recA1*, *endA1*, *thi*, *gyrA96*, *supE44*, *hsdR17*Δ (*lac-proAB*) /F’[*traD36,proAB*$^{+}$, *lacI*$^{q}$, *lacZΔ M15*] | Laboratory stock |
| *B. subtilis* | Wild-type *Bacillus subtilis* 168 | Laboratory stock |
| BS-P43-sfGFP | Wild-type *B. subtilis* carrying the plasmid PXY-P43-sfGFP | Laboratory stock |
| BS-A1-sfGFP | Wild-type *B. subtilis* carrying the plasmid PXY-A1-sfGFP | This study |
| BS-A2-sfGFP | Wild-type *B. subtilis* carrying the plasmid PXY-A2-sfGFP | This study |
| BS-A3-sfGFP | Wild-type *B. subtilis* carrying the plasmid PXY-A3-sfGFP | This study |
| BS-A4-sfGFP | Wild-type *B. subtilis* carrying the plasmid PXY-A4-sfGFP | This study |
| BS-A5-sfGFP | Wild-type *B. subtilis* carrying the plasmid PXY-A5-sfGFP | This study |
| BS-A6-sfGFP | Wild-type *B. subtilis* carrying the plasmid PXY-A6-sfGFP | This study |
| BS-A7-sfGFP | Wild-type *B. subtilis* carrying the plasmid PXY-A7-sfGFP | This study |
| BS-A8-sfGFP | Wild-type *B. subtilis* carrying the plasmid PXY-A8-sfGFP | This study |
| BS-A9-sfGFP | Wild-type *B. subtilis* carrying the plasmid PXY-A9-sfGFP | This study |
| BS-A10-sfGFP | Wild-type *B. subtilis* carrying the plasmid PXY-A10-sfGFP | This study |
| BS-A11-sfGFP | Wild-type *B. subtilis* carrying the plasmid PXY-A11-sfGFP | This study |
| BS-A12-sfGFP | Wild-type *B. subtilis* carrying the plasmid PXY-A12-sfGFP | This study |
| BS-A13-sfGFP | Wild-type *B. subtilis* carrying the plasmid PXY-A13-sfGFP | This study |

| | | |
|---|---|---|
| BS-A14-sfGFP | Wild-type *B. subtilis* carrying the plasmid PXY-A14-sfGFP | This study |
| BS-A15-sfGFP | Wild-type *B. subtilis* carrying the plasmid PXY-A15-sfGFP | This study |
| BS-A16-sfGFP | Wild-type *B. subtilis* carrying the plasmid PXY-A16-sfGFP | This study |
| BS-A17-sfGFP | Wild-type *B. subtilis* carrying the plasmid PXY-A17-sfGFP | This study |
| BS-A18-sfGFP | Wild-type *B. subtilis* carrying the plasmid PXY-A18-sfGFP | This study |
| BS-A19-sfGFP | Wild-type *B. subtilis* carrying the plasmid PXY-A19-sfGFP | This study |
| BS-A20-sfGFP | Wild-type *B. subtilis* carrying the plasmid PXY-A20-sfGFP | This study |
| BS-A21-sfGFP | Wild-type *B. subtilis* carrying the plasmid PXY-A21-sfGFP | This study |
| BS-A22-sfGFP | Wild-type *B. subtilis* carrying the plasmid PXY-A22-sfGFP | This study |
| BS-PlepA-sfGFP | Wild-type *B. subtilis* carrying the plasmid PXY- PlepA-sfGFP | This study |
| BS-Pveg-sfGFP | Wild-type *B. subtilis* carrying the plasmid PXY- Pveg-sfGFP | This study |
| JM109-A1-sfGFP | Wild-type JM109 carrying the plasmid PXY-A1-sfGFP | This study |
| JM109-A2-sfGFP | Wild-type JM109 carrying the plasmid PXY-A2-sfGFP | This study |
| JM109-A3-sfGFP | Wild-type JM109 carrying the plasmid PXY-A3-sfGFP | This study |
| JM109-A4-sfGFP | Wild-type JM109 carrying the plasmid PXY-A4-sfGFP | This study |
| JM109-A5-sfGFP | Wild-type JM109 carrying the plasmid PXY-A5-sfGFP | This study |
| JM109-A6-sfGFP | Wild-type JM109 carrying the plasmid PXY-A6-sfGFP | This study |
| JM109-A7-sfGFP | Wild-type JM109 carrying the plasmid PXY-A7-sfGFP | This study |
| JM109-A8-sfGFP | Wild-type JM109 carrying the plasmid PXY-A8-sfGFP | This study |
| JM109-A9-sfGFP | Wild-type JM109 carrying the plasmid PXY-A9-sfGFP | This study |
| JM109-A10-sfGFP | Wild-type JM109 carrying the plasmid PXY-A10-sfGFP | This study |
| JM109-A11-sfGFP | Wild-type JM109 carrying the plasmid PXY-A11-sfGFP | This study |
| JM109-A12-sfGFP | Wild-type JM109 carrying the plasmid PXY-A12-sfGFP | This study |
| JM109-A13-sfGFP | Wild-type JM109 carrying the plasmid PXY-A13-sfGFP | This study |
| JM109-A14-sfGFP | Wild-type JM109 carrying the plasmid PXY-A14-sfGFP | This study |
| JM109-A15-sfGFP | Wild-type JM109 carrying the plasmid PXY-A15-sfGFP | This study |
| JM109-A16-sfGFP | Wild-type JM109 carrying the plasmid PXY-A16-sfGFP | This study |
| JM109-A17-sfGFP | Wild-type JM109 carrying the plasmid PXY-A17-sfGFP | This study |
| JM109-A18-sfGFP | Wild-type JM109 carrying the plasmid PXY-A18-sfGFP | This study |
| JM109-A19-sfGFP | Wild-type JM109 carrying the plasmid PXY-A19-sfGFP | This study |
| JM109-A20-sfGFP | Wild-type JM109 carrying the plasmid PXY-A20-sfGFP | This study |
| JM109-A21-sfGFP | Wild-type JM109 carrying the plasmid PXY-A21-sfGFP | This study |

| | | |
|---|---|---|
| JM109-A22-sfGFP | Wild-type JM109 carrying the plasmid PXY-A22-sfGFP | This study |
| JM109-Ran1-sfGFP | Wild-type JM109 carrying the plasmid PXY- Ran1-sfGFP | This study |
| JM109-Ran2-sfGFP | Wild-type JM109 carrying the plasmid PXY- Ran2-sfGFP | This study |
| JM109-Ran3-sfGFP | Wild-type JM109 carrying the plasmid PXY- Ran3-sfGFP | This study |
| JM109-J23100-sfGFP | Wild-type JM109 carrying the plasmid PXY- Ran1-sfGFP | This study |
| JM109-J23102-sfGFP | Wild-type JM109 carrying the plasmid PXY- Ran2-sfGFP | This study |
| JM109-J23119-sfGFP | Wild-type JM109 carrying the plasmid PXY- Ran3-sfGFP | This study |

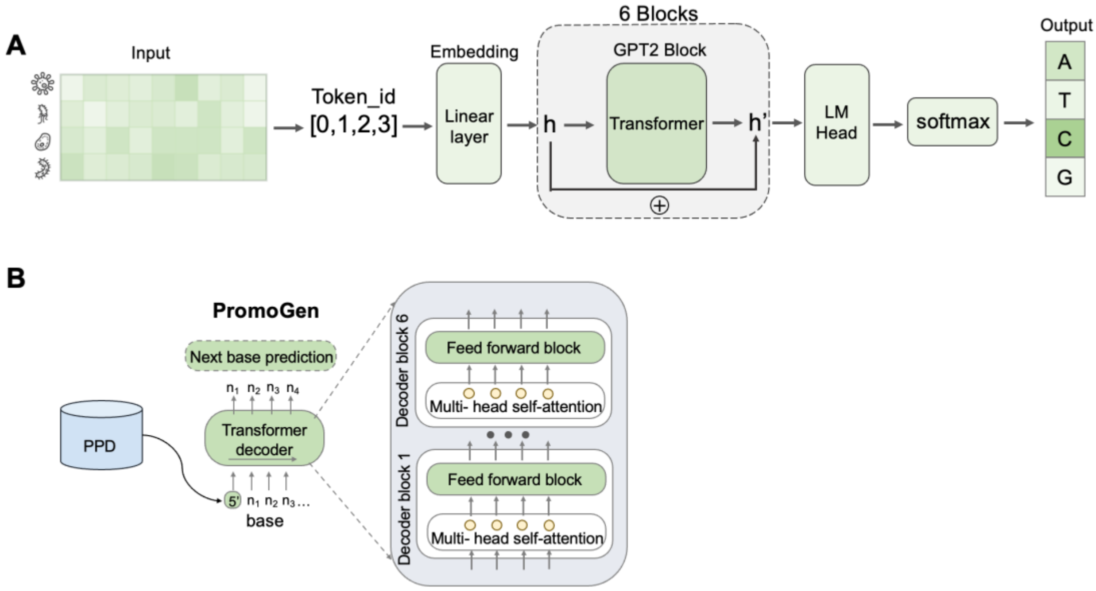


Figure S1. The model architecture of PromoGen. (A) Transformer architecture and objections of this work. Input is the promoters from various species, and the output is the probability of A, T, C, and G. (B) Illustration of the PromoGen.

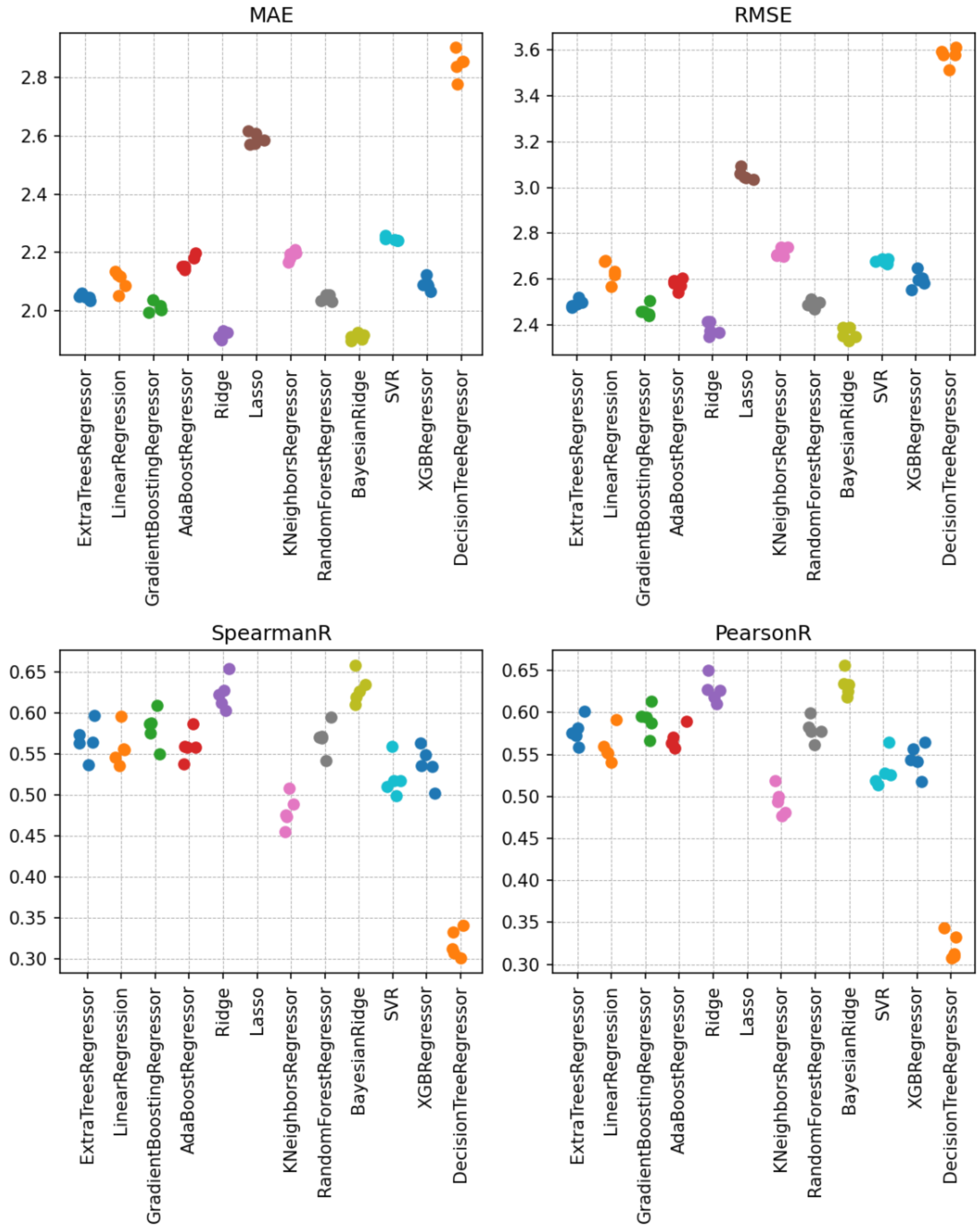


Figure S2. The comparison involved 12 predictive models, with the primary evaluation metrics encompassing Mean Absolute Error (MAE), Root Mean Squared Error (RMSE), Spearman correlation coefficient, and Pearson correlation coefficient.

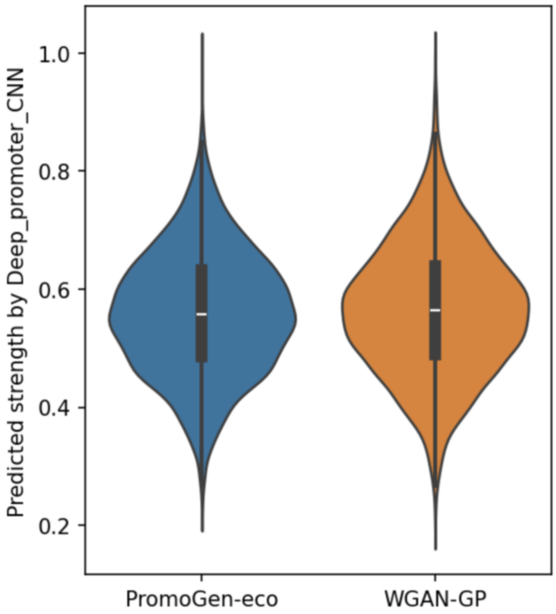


Figure S3. The comparative analysis between PromoGen-eco and WGAN-GP was conducted using a Convolutional Neural Network (CNN) predictive model as reported by Wang.

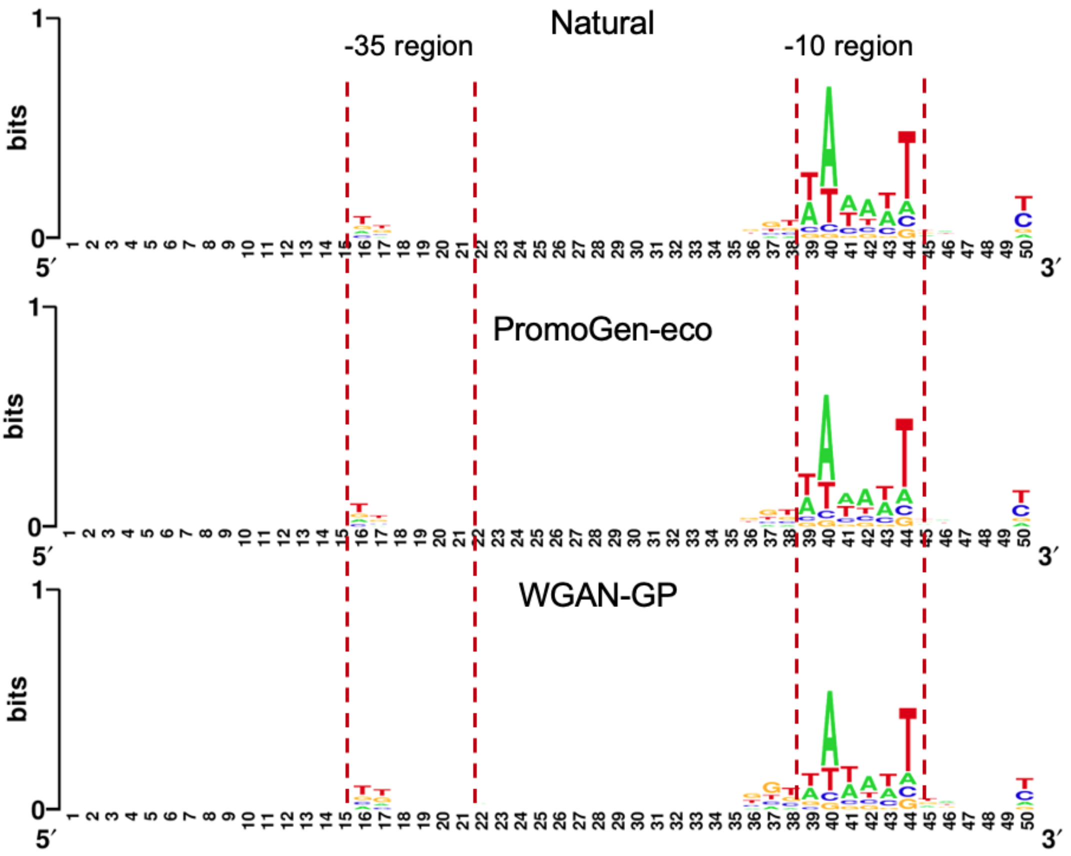


Figure S4. The sequence logos of the PromoGen-eco generated, native, and WGAN-GP- generated promoters.

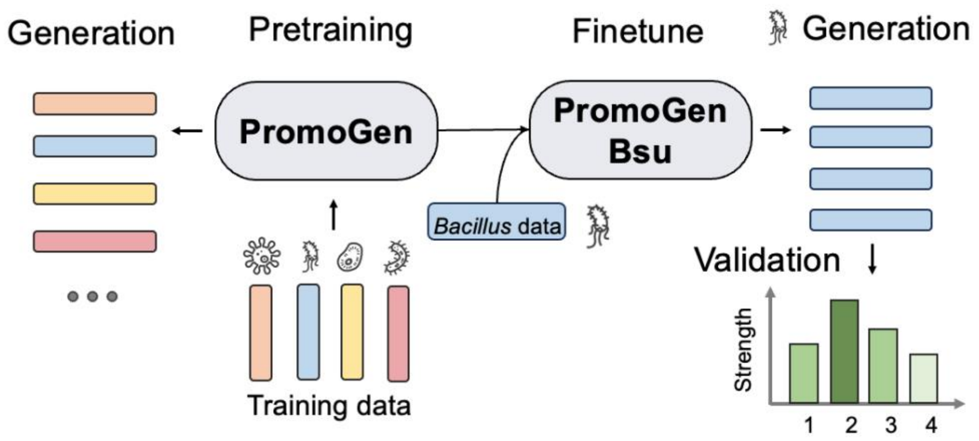


Figure S5. The PromoGen-pre was first pretrained on larger dataset containing promoter sequences from multiple species. To generate promoter that will be functional in a particular host, *Bacillus* for example, the PromoGen-pre model is further finetuned using a small amount of data containing only *Bacillus* promoters, the finetuned version is termed PromoGen-bsu. The sequences generated by PromoGen-bsu is subjected to experimental verification.

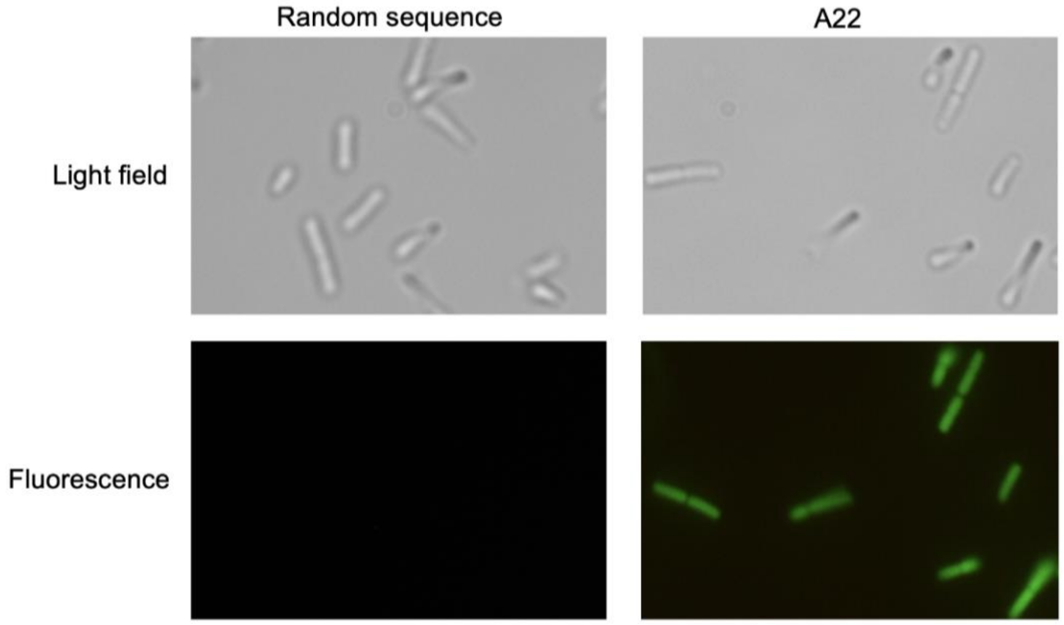


Figure S6. The fluorescence of *B. subtilis* expressing the sfGFP under the random sequence and A22.

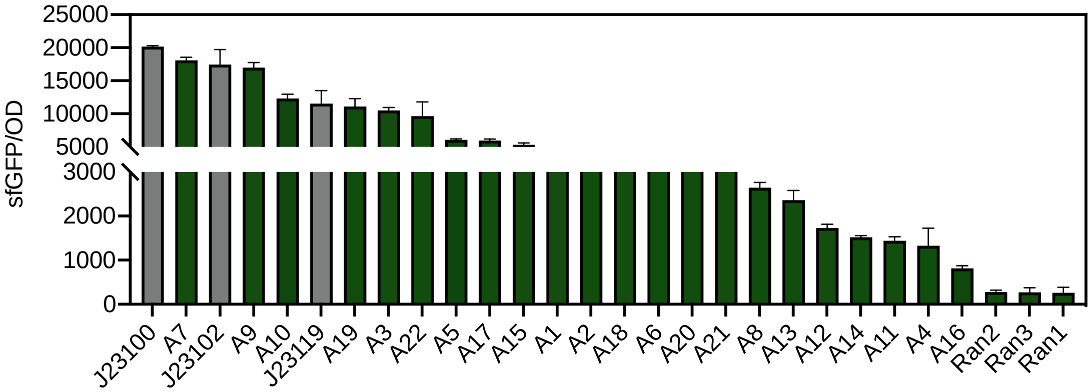


Figure S7. The validation of promoter strength was confirmed by measuring the fluorescence intensity of sfGFP in *E. coli*. The experiment utilized three robust promoters, namely J23100, J23102, and J23119, as positive controls, while three randomly generated sequences served as negative controls.
